# Substrate stiffness modulates integrin α5 expression and ECM-associated gene expression in fibroblasts

**DOI:** 10.1101/2021.11.22.469526

**Authors:** Brijesh Kumar Verma, Aritra Chatterjee, Paturu Kondaiah, Namrata Gundiah

**Author notes:** For correspondence: **Paturu Kondaiah**, Department of Molecular Reproduction, Development, and Genetics, Indian Institute of Science, Bangalore 560012. **Email**, **Namrata Gundiah**, Biomechanics Laboratory, Department of Mechanical Engineering, Indian Institute of Science, Bangalore 560012. **Email**, **Tel (Off/ Lab)**: 91 80 2293 2860/ 3366.

## Abstract

Biomaterials, like polydimethylsiloxane (PDMS), are soft, biocompatible, and tuneable, which makes them useful to delineate specific substrate factors that regulate the complex landscape of cell-substrate interactions. We used a commercial formulation of PDMS to fabricate substrates with moduli 40 kPa, 300 kPa, and 1.5 MPa, and cultured HMF3S fibroblasts on them. Gene expression analysis was performed by RT-PCR and Western blotting. Cellular and nuclear morphologies were analyzed using confocal imaging, and MMP-2 and MMP-9 activities were determined with gelatin zymography. Results, comparing mechanotransduction on PDMS substrates with control petridishes, show that substrate stiffness modulates cell morphologies and proliferations. Cell nuclei were rounded on compliant substrates and correlated with increased tubulin expression. Proliferations were higher on stiffer substrates with cell cycle arrest on softer substrates. Integrin α5 expression decreased on lower stiffness substrates, and correlated with inefficient TGF-β activation. Changes to the activated state of the fibroblast on higher stiffness substrates were linked to altered TGF-β secretion. Collagen I, collagen III, and MMP-2 expression levels were lower on compliant PDMS substrates as compared to stiffer ones; there was little MMP-9 activity on substrates. These results demonstrate critical feedback mechanisms between substrate stiffness and ECM regulation by fibroblasts which is highly relevant in diseases like tissue fibrosis.

## 1. Introduction

Cells are exquisitely sensitive to their milieu and modulate behaviours in response to physical and biochemical cues from the microenvironment [1-3]. Fibrous collagen and elastin networks, in combination with proteins such as fibronectin and laminin, help attach cells to the extracellular matrix (ECM) through integrin receptors in the cell membrane [4]. These connections play a critical role in directing cell shape, proliferation, differentiation, migration, and secretion of biochemical molecules [5]. Tissue fibrosis is a result of chronic inflammation, initiated due to the immune response, and accompanied with the recruitment of growth factors, proteolytic enzymes, cytokines, and angiogenic factors secreted by transformed fibroblasts that progressively remodel the affected extracellular matrix (ECM) [6, 7]. Fibrosis occurs when the synthesis of *de novo* collagen by fibroblasts exceeds the rate at which matrix gets degraded by matrix metalloproteinases (MMPs), and is a primary reason underlying the dysfunction of tissues like myocardium after infarction, aneurysm growth and subsequent rupture, and in organs such as liver, bladder and lungs [6-8].

Complex and intricate feedback processes between cellular tension and the extracellular physical properties influence the hierarchical assembly and ECM microstructure during remodeling [5,9]. Mechanosensory, heterodimeric, transmembrane integrins link the internal cytoskeleton of cells to the ECM, and participate in the deposition of ECM proteins, including thrombospondin and collagen type I [9]. Collagens perform the primary function of load bearing, provide strength to tissues, and help transfer stresses reversibly during deformation to the attached cells. ECM moduli are highly variable across different organs and range from ∼100 Pa in the brain and fat tissues, to about 10 kPa for muscle, and orders of magnitude higher in arteries and the cartilage [10]. Substrate stiffness is a critical regulator of fibroblast morphology; cell stiffness is tuned to the underlying substrate stiffness [10]. Embryonic stem cells, cultured on polydimethylsiloxane (PDMS) substrates of varied stiffness, show varied differentiation, spreading, and proliferation responses [11]. Collagen hydrogel networks, with thick fibers, large pores, and higher moduli, promote mechanosignaling in stromal cells, and their consequent differentiation into myofibroblasts [12]. The tissue stiffness undergoes dramatic change during fibrosis due to the differential expression of ECM proteins, their subsequent cross-linking, and that of proteases, including matrix metalloproteinases (MMP’s). Together, these alter the physical properties of the ECM, and biochemically influence signaling through the release of small bioactive peptides and growth factors [8, 9].

Biomaterials are useful to delineate the role of specific substrate factors that regulate the complex landscape of cell-substrate interactions. Commonly used materials for mechanotransduction include biologically derived scaffolds, such as Matrigel, collagen I, fibrin, and synthetic polymers such as poly acrylamide and gelatin methacrylate gels (GelMA) [14, 15]. Natural biomaterials are attractive because they recapitulate the *in vivo* substrate characteristics but have significant challenges due to biological variability, degradability, variability in pore sizes, and other factors that make data interpretation difficult. In contrast, synthetic biomaterials, with surfaces modified to provide motifs for cell attachment, are attractive as they can be repeatably tuned, and permit control of mechanical characteristics such as modulus, surface topography, and pore size.

We test the sensitivity and specificity of human mammary fibroblast (HMF3S) responses to changes in substrate stiffness using a well characterized soft, biocompatible, and tuneable PDMS elastomer. We select substrate properties to broadly represents collagenous matrix/ muscle tissue (∼40 kPa), collagenous bone (∼300 kPa), cartilage (∼1500 kPa), and bone (∼10^7^ kPa) respectively [16]. We explore the role of material modulus on integrin α5, collagen (types I and III), and MMP (2 and 9) expressions. Our results clearly demonstrate the regulation and feedback mechanisms between cell interactions with the underlying substrate properties. Cell proliferation was also crucially dependent on substrate stiffness. Most current treatments for fibrotic diseases are aimed at targeting the inflammatory response to slow the progression of fibrosis or partially reverse effects to restore homeostasis [3]. Investigations into the mechanobiology of fibroblast interactions with the ECM are critical, but challenging, in characterizing the signaling pathways that drive MMP secretion and collagen deposition in the evolving fibrous tissue.

## 2. Materials and methods

### 2.1. Cell lines

HMF3Ss cells were a generous gift from Dr. P. Jat at the Ludwig Cancer Institute, UK. HMF3Ss, HT1080 (fibrosarcoma) and CCL-64 PAI cells were propagated in DMEM containing 10% fetal calf serum in 95% humidified CO_2_ incubator maintained at 37°C in the study.

### 2.2. Preparation of PDMS substrates with various substrate stiffness for cell culture

Glass coverslips were cleaned by ultrasonication in sterile distilled water, 1 mM EDTA, 70% ethanol, and 100% ethanol for 15 minutes each, and air dried. Substrates of different stiffness were prepared by mixing silicone elastomers (Sylgard® 184, Dow Corning) using the cross-linker in different stoichiometric weight ratios of 10:1 (1.58±0.188 MPa), 20:1 (351±24 kPa), and 40:1 (41.89±4.09 kPa) based on protocols developed in the laboratory [16]. The solution was degassed in a desiccator to remove bubbles, the mixture spin coated on cleaned coverslips at 300 rpm for 2 minutes, and the materials cured in an oven at 80°C for 2 hours. Crosslinked PDMS substrates were plasma treated for 2 minutes, and incubated with 40 μg/ml of fibronectin for 1 hour at 37°C. The substrates were washed twice with PBS and cultured using HMF3S cells with 8000-10000 cells/cm^2^cell density.

### 2.3. RNA isolation and Real-time PCR

RNA was isolated using TRI™ reagent (Sigma-Aldrich, USA) based on protocols suggested by the manufacturer. Briefly, cells were washed in Phosphate Buffered Saline (PBS; pH 7.0), lysed using TRI reagent at room temperature, and chloroform added to the mixture to separate the solution phase. The solution was centrifuged at 12000g for 10 minutes, and RNA from the cells was precipitated in the aqueous phase using 50% isopropanol. The resultant was suspended in 70% ethanol, and the RNA dissolved in 20 μl RNase-free water. Purity and integrity of RNA were tested using the ratio of absorbances at 260 and 280 nm, and formaldehyde agarose gels. A cDNA synthesis kit (Applied Biosystems, USA) was used to reverse transcribe 2 μg RNA based on protocols recommended by the manufacturer. Gene specific primers were used to test the expression levels of various proteins of interest. GAPDH (Glyceraldehyde 3 phosphate Dehydrogenase) was used as a normalizing control. A list of primers used in this study is included as supplementary table S1.

### 2.4. Western blot Analysis

Cells were washed with PBS and lysed in RIPA (Radio-immunoprecipitation assay) buffer containing 50 mM Tris-HCl buffer (pH 7.4), 1% NP-40, 0.25% deoxycholate, 150 mM NaCl, 1 mM EDTA (pH 8.0), 1 mM PMSF, 1 mM NaF, 1 mM sodium orthovanadate, 1x protease inhibitor cocktail III (stock 100x, Calbiochem, USA), and 0.1% SDS. Lysates were stored in ice for 10-15 minutes and centrifuged for 10 minutes at 12,000g and at 4°C. 20 μg of the total cell lysate was resolved by 12.5% SDS-PAGE and blotted onto immobilon membranes by electroblotting. Antibodies used in this study include pFAK Tyr397 (#3283S, CST, USA), FAK (#3285, CST, USA), ITGA5 (#ITT05200, GTI, USA), tubulin α (#ITT4777H, GTI, USA), β-actin (#ITM2093, GTI, USA), and GAPDH (#3683S, CST, USA).

### 2.5. Immunocytochemistry

Cells were fixed using 3.7% paraformaldehyde (PFA) for 25 minutes and blocked with 10% FBS at room temperature for 1 hour. To visualize the actin cytoskeleton, cells were incubated with rhodamine phalloidin (Thermofisher, 1:200) for 60 minutes. Hoechst stain (Thermofisher, 1:400) for 15-20 minutes was used for the nucleus. Samples were imaged using a confocal microscope (Airy Scan ZEISS LSM 900 with Airyscan 2) at 10X, and 63X magnifications, respectively.

### 2.6. Zymography

Zymography was performed to examine the enzymatic activity of matrix metalloproteinases (MMP-2 and MMP-9) in the conditioned media of HMF3S cells. In this method, cells were seeded on PDMS substrates of different stiffness, the conditioned medium collected, and centrifuged (5000 g) to pellet the cell debris. The spent medium was transferred to a fresh tube and 20 µl of conditioned media was loaded on a 12.5% SDS-PAGE gel with 0.5% gelatin for electrophoresis. In addition to the SDS-PAGE - gelatin gel, the conditioned media was also resolved on one normal 12.5% SDS-PAGE. Normal gel was used as loading control. The resulting gels were incubated in Triton X 100 (2.5%) for 30 - 40 minutes and placed overnight in an incubation buffer (Tris-HCl buffer pH 7.5-10mM, Triton X 100-1.25%, CaCl_2_-5mM, ZnCl_2_-1µM; 37^0^C) to quantify the enzymatic activity. Coomassie brilliant blue (15-20 minutes at room temperature) was used to stain the gels. Images of the destained gel, obtained using methanol (40%), glacial acetic (10%), and water (50%), were obtained using a Chemidoc imaging system (ChemiDoc XRS and Gel Imaging System).

### 2.7. Cell cycle analysis

HMF3S cells were seeded on the different substrates for 24 hours, washed with chilled PBS, and lysed for 30 minutes using iced hypotonic solution containing propidium iodide (PI), sodium citrate, Triton X-100, and double-distilled water. The cell lysate was centrifuged at 600g for 5 minutes at 4°C, the supernatant discarded, and the pellet containing the nuclei resuspended in hypotonic solution containing propidium iodide (PI). Samples were analysed using FACS (BD Accuri C6).

### 2.8. Trypan blue exclusion assay

Cells were seeded in 35 mm dishes for 24 hours, trypsinized, and incubated in 0.4% trypan blue solution for 3 minutes at room temperature. Live cells were counted using a haemocytometer.

### 2.9. Measurement of cell and nuclear morphometrics

The cell and nuclear aspect ratios were calculated using ImageJ (NIH) software [17]. Images of single cells, obtained using confocal imaging, were segmented to identify the cell boundaries, and the corresponding areas were obtained from the binarized images. The nucleus was fit to an ellipse and the aspect ratio determined as a ratio of the major to the minor axis [18].

### 2.10. Statistical analysis

Statistical differences in the mean values between groups, using three biological repeats, were obtained using ANOVA with Bonferroni multiple comparison tests in GraphPad Prism (v6.0, GraphPad Software, Inc., La Jolla, CA, USA). p-values < 0.05 were considered statistically significant.

## 3. Results and discussion

### 3.1. Substrate stiffness modulates fibroblast morphology and proliferation

We used PDMS of various stiffness ratios to represent the native tissue stiffness in this study and quantified the expression levels of cytoskeletal proteins, actin and tubulin, on the various substrates in this study. Results from cells cultured on petri dish substrates were included as controls (Fig. 1a). We compared the optical densities of proteins on the various substrates relative to GAPDH levels (n=3). Fig. 1b shows that actin decreased on substrates with lower stiffness; significant results were discernible between control and the 40:1 PDMS substrates alone (p<0.05). In contrast, tubulin significantly increased with decrease in substrate stiffness (Fig. 1c). The differences in tubulin expression levels are apparent on the 20:1 and 40:1 PDMS substrates as compared to the other substrate stiffnesses in this study.

**Fig. 1:**
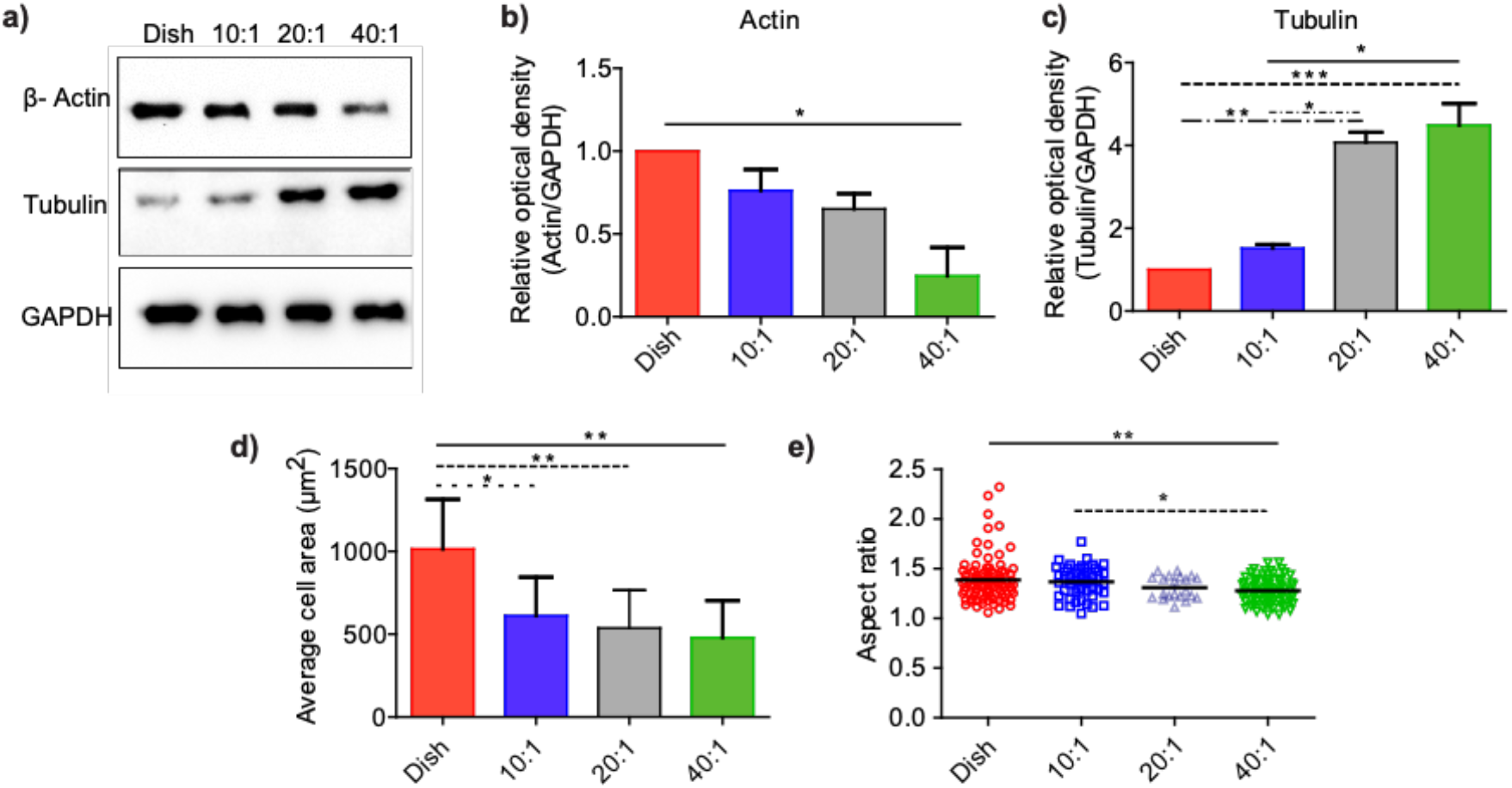
Substrate stiffness regulates the cytoskeletal gene expression. **(a)** Immunoblot of β-actin and tubulin from cell lysate collected from cell seeded on dish, 10:1, 20:1 and 40:1 substrate. Densitometric quantitation using ImageJ were obtained for (**b**) β-actin and (**c**) tubulin using the immunoblots. (**d**) Average fibroblast areas on substrates of varied stiffness were obtained using the confocal images (N∼20 for each group). **(e)**. Aspect ratios of cells on substrates of varied stiffness were obtained using analysis from confocal image of nucleus (N∼40 for each group). All the experiments were performed in biological triplicates. Data are represented as mean ± SEM; * p<0.05; ** p<0.01; *** p<0.001.

Poly(dimethylsiloxane) has many properties, including gas permeability, biocompatibility, low water absorption, ease of microfabrication, high level of stability, and mechanical property tuneability which make it a favorable biomaterial in mechanobiology experiments [17,20,21]. The inherent intrinsic hydrophobicity of PDMS dictates interactions with biological samples that adhere to the surface; there are however few differences in the amount of adsorbed protein to the substrates [22,23]. Plasma treatment of PDMS results in a hydrophilic and rough surface through surface oxidization [24]. The adhesion of cells to substrates is dictated by several factors such as nature of the ligand coating, substrate stiffness, and cell types; there are however no clear correlations between cell morphologies and the underlying substrate stiffness [25]. The adherent fibroblast areas on PDMS substrates in our study show significant differences as compared to cells on control petridishes (Fig. 1d; Fig. S1). The average cell spread area (n > 40 for each group) was lower on PDMS substrates with lower stiffness as compared to those cultured on control petridishes (p<0.01). There were however no differences in the cell spread areas on PDMS substrates of various stiffness in this study. Nuclei on stiffer substrates were less rounded on control substrates as compared to those on compliant substrates. The aspect ratios of nuclei decreased with a decrease in substrate stiffness; these data clearly demonstrate a rounding of the nucleus on compliant substrates (Fig. 1e).

Because microtubules are the primary components of the mitotic spindle, we hypothesized that cell proliferations should differ on substrates of differing moduli. Fig. 2a shows differences in the cell growth over a 3 day period on PDMS substrates of varied stiffness. Cellular growth on stiffer substrates over the 24 hour period was significantly higher on the control substrate as compared to the (40:1) compliant PDMS substrate (p<0.01). Although cells proliferated on all PDMS substrates, those seeded on higher stiffness substrates showed a higher rate of proliferation at each of the three different time periods.

**Fig. 2:**
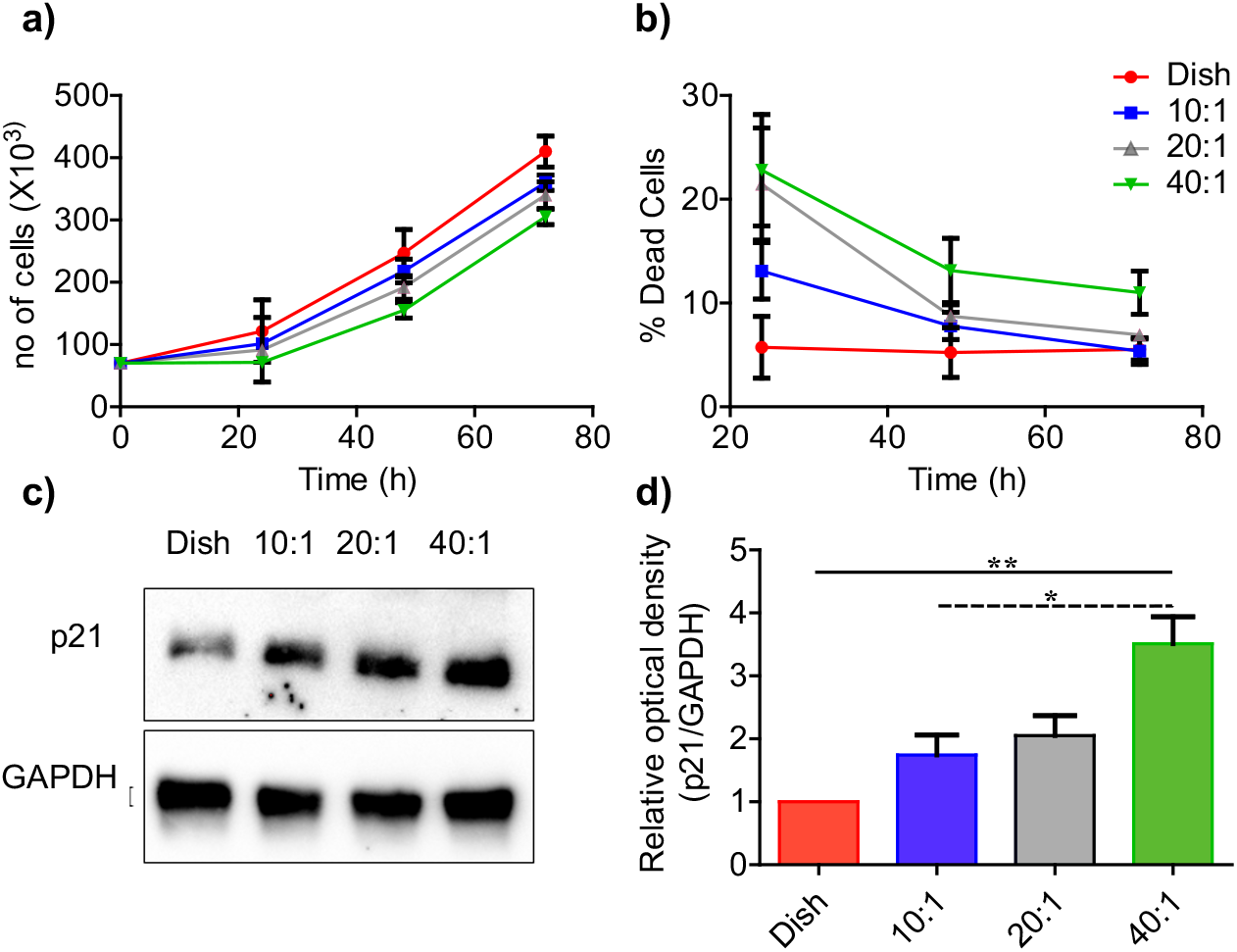
Reduction in substrate stiffness restricts cell growth. **(a)** Cell proliferations on petridish, 10:1, 20:1, and 40:1 PDMS substrates were obtained using trypan blue assay. (**b)** The percentage of dead cells in each group are shown at 24 hours, 48 hours, and 72 hours. **(c)** The immunoblot of p21, and (**d**) densitometric analysis of immunoblot using ImageJ. All experiments are performed in biological triplicates. Data are represented as mean ± SEM; * p<0.05; ** p<0.01.

We used the trypan blue live-dead cell assay to distinguish between dead and live cells on all substrates in the study. Control substrates had < 10% dead cells at 24 hours in contrast to (10:1, 20:1 and 40:1) PDMS substrates that showed over 10% cell death (Fig. 2b). The percent dead cells in the population decreased significantly at later time points. All substrates, barring the 40:1 substrate, had < 10% dead population at 72 hours. We used FACS base cell cycle profiling for cells cultured on substrates of different stiffness. Cells were initially synchronised through serum starvation for 24 hours and normal media was added to release the cell cycle. Samples were collected and assayed using FACS (Fig. S2). Cells were enriched in G0/G1 cycle phase on the lower stiffness PDMS substrates which suggests possible cell cycle arrest. We next quantified the expression of p21 protein as a marker for cell cycle arrest [25]. Results show that p21 protein expression was significantly higher on lower stiffness substrates as compared to control substrates; these results corroborate the effects of substrate stiffness on cell cycle progression.

These data agree with others that demonstrate increased proliferation of mammary epithelial cells, vascular smooth muscle cells, and mouse embryonic fibroblasts on higher stiffness substrates fabricated using alginate [25, 27]. Increased focal adhesion kinase (FAK) activity on stiffer substrates is linked to the higher cell proliferations [28]. Studies also suggest the existence of a possible tuning mechanism in cells to substrates of stiffness similar to the target tissue from which they are derived [10]. The specific mechanisms underlying these behaviors are presently not well understood.

### 3.2. Regulation of integrin α5 expression with substrate stiffness

Integrins promote the spreading of cells on substrates and participate in signaling through cross-talk between the substrate ligands and the cell cytoskeleton. We tested the effect of substrate stiffness on integrin α5 expression in HMF3S cells cultured on PDMS substrates of different stiffness, and used the culture dish as control. Integrin α5 domains in fibroblasts selectively attach through fibrillar adhesions to fibronectin ligands on the substrate. These interactions change the state of the integrins from low to high affinity conformations and help anchor the cell to the substrate [29]. Signaling adaptors of integrins include kinases, like FAK, that are highly dynamic and recruit other molecules to help organize the actin cytoskeleton [29]. We studied changes in the expression levels of integrin α5 (ITGA5), FAK and phospho-FAK to test for differences in cell signaling on varying stiffness PDMS substrates. Fig. 3a shows a substantial reduction in ITGA5 and phospho-FAK proteins on PDMS substrates (20:1 and 40:1) as compared to the dish. We next compared mRNA expression of ITGA5 gene by qRT-PCR; results show clear differences between the control and PDMS substrates (p<0.05; Fig. 3b). FAK tyr397 phosphorylation showed a direct correlation with the integrin α5 protein expression on these substrates (Fig. 3a). These results suggest that changes to the substrate stiffness regulates the integrin α5 expression on PDMS substrates and also alters cell signaling through activation of FAK. Mechanically induced changes to the conformation of integrins may hence help in mechanotransduction to sense the substrate characteristics.

**Fig. 3:**
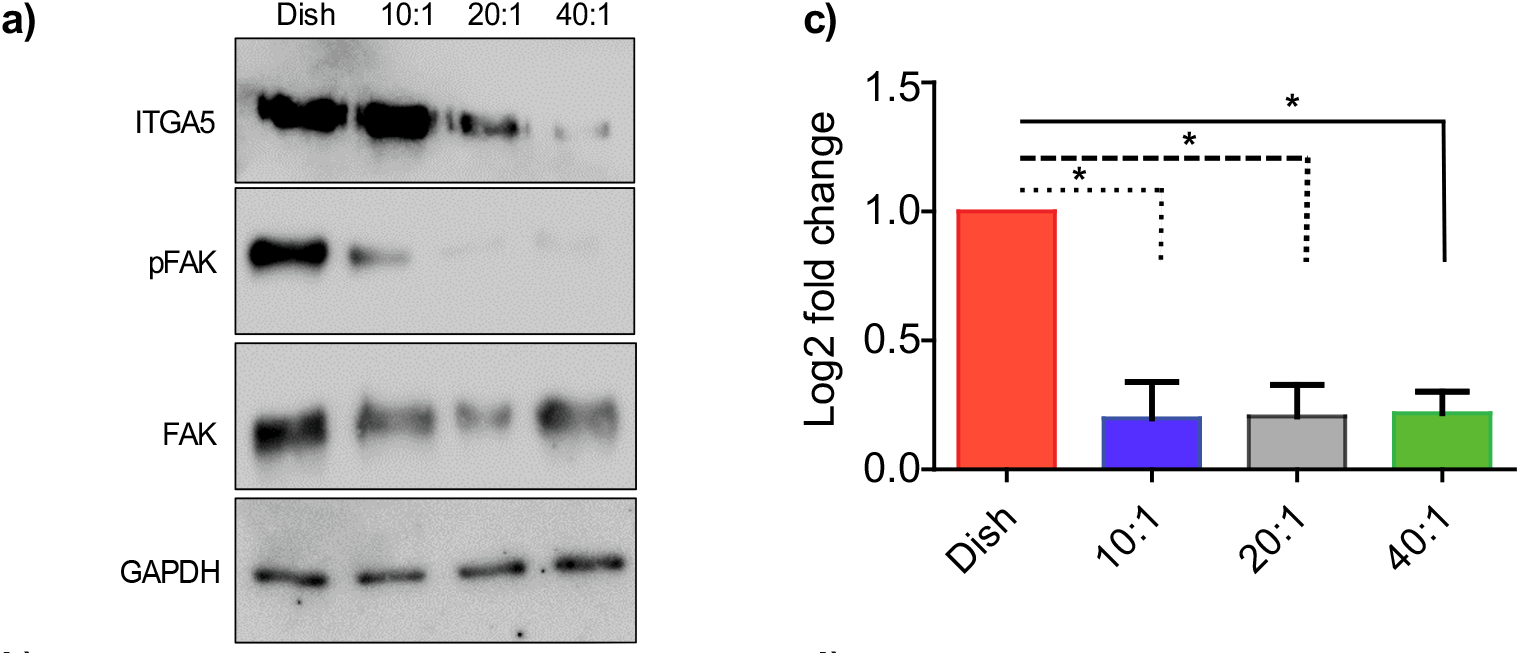
Substrate stiffness regulates the expression of integrin α5. **(a)** Immunoblots of integrin α5, FAK and pFAK from cell lysates collected from cells seeded on dish, 10:1, 20:1 and 40:1 PDMS substrates respectively. **(b)** mRNA expression of integrin α5 is shown from four different conditions. * p<0.05; ** p<0.01

### 3.3. ECM-associated genes are downregulated at lower stiffness

We quantified differences in the expression levels of collagen I, collagen III and MMP’s on the PDMS substrates of differential stiffness and compared results from these experiments using fibroblasts on control petri dishes. Fig. 4a shows that the Col I mRNA expression gradually decreased with a decrease in PDMS substrate stiffness. There were also significant differences between the control substrate and prepared PDMS substrates of different stiffness. We used Real time PCR to analyse the expressions of collagen I, collagen III and MMP genes in cells cultured on substrates with different stiffnesses. The expression levels of these genes were significantly lower on compliant substrates as compared to stiffer ones (Fig. 4a and 4b). We hypothesized that the substrate stiffness may also affect MMP activity. We used gelatin zymography to assess the effects of substrate stiffness on MMP-2 and MMP-9 activities (Fig. 5a). The condition media of cells cultured on different substrate stiffnesses was resolved on 12.5% of SDS-PAGE containing 5% gelatin. The gel was subsequently transferred to the MMP activation buffer. Regions of gelatin, degraded from gel due to MMP activity, were not stained with Coomassie Blue. HT1080 cell condition media, containing both MMP-2 and MMP-9, were used as positive control. The conversion of MMP-2 to active-MMP-2, using gelatin zymography, was significantly higher on lower stiffness substrates (Fig. 5a). In addition, the mRNA expression of MMP2 gene was higher on lower stiffness as compared to that on higher stiffness (Fig. 5b). Intriguingly, there was no significant activity of MMP-9 in the conditioned media (Fig. 5a). Our results clearly demonstrate that HMF3S fibroblast cells alter their matrix production (Col I and Col III) and MMP-2 secretion based on the stiffness of their microenvironment. Lachowski and co-workers showed that the modulus of polyacrylamide substrates modulates gene expressions of MMP-2, MMP-9, and TIMP-1 in hepatic stellate cells [30]. The moduli of substrates investigated in their study varied from 4 kPa to 25 kPa which is significantly lower and over a narrower range as compared to that in our study.

**Fig. 4:**
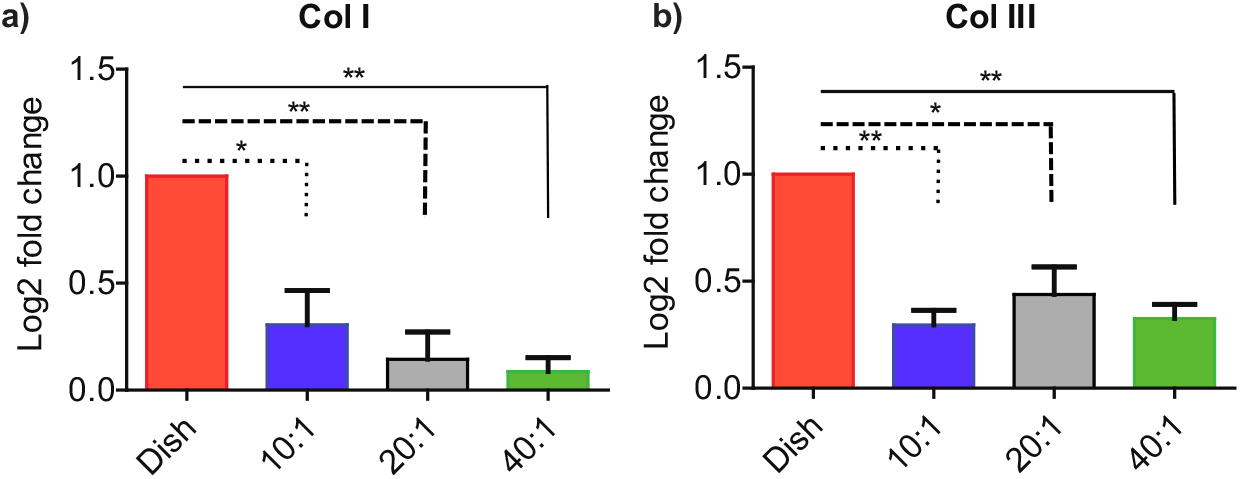
Variations in ECM-associated gene expression and activity with substrate stiffness. qRT-PCR expression analysis of (**a**) Col I, and (**b**) Col III are shown from RNA collected from cells cultured on dish, 10:1, 20:1, and 40:1 substrates respectively. All experiments were performed in biological triplicates. Data are represented as mean ± SEM; * p<0.05; ** p<0.01.

**Fig. 5:**
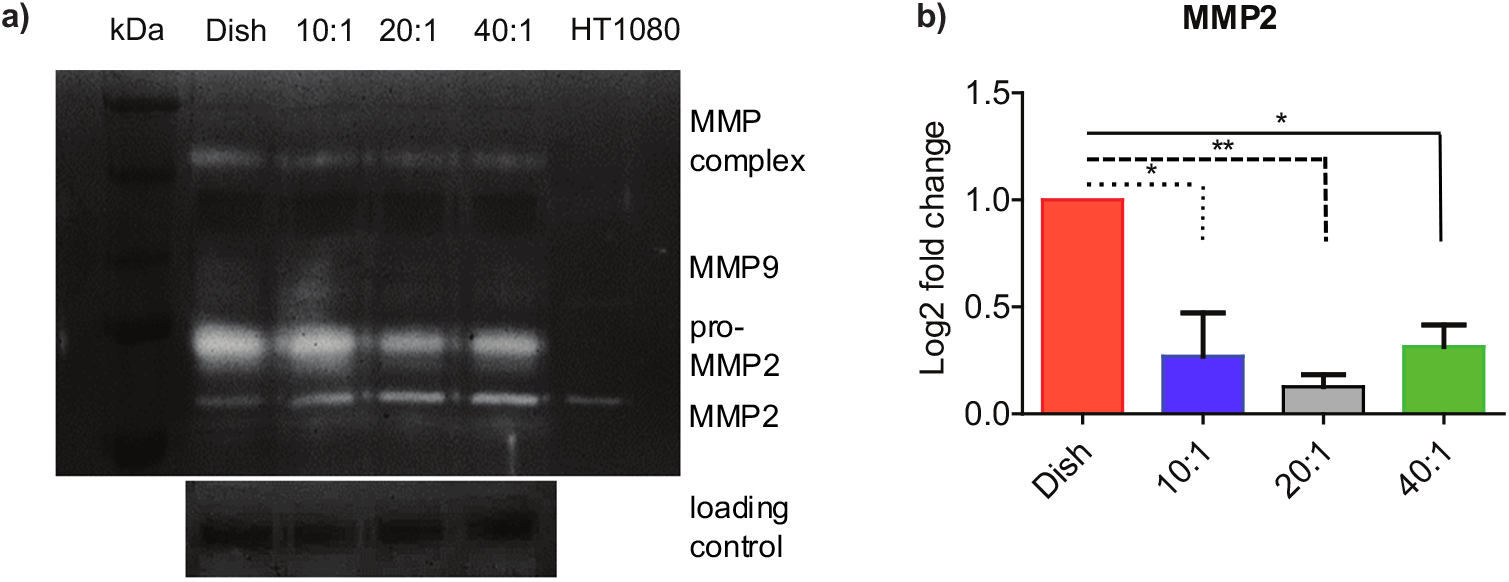
MMP activity depends on the substrate stiffness. **(a)** Gelatin zymograph with condition media collected from cells cultured on dish, 10:1, 20:1 and 40:1 PDMS substrates were assayed for MMP-2 and MMP-9. (**b**) qRT-PCR expression analysis of MMP-2 from RNA collected from cells cultured on substrates with variable stiffnesses. All experiments were performed in biological triplicates. Data are represented as mean ± SEM; * p<0.05; ** p<0.01.

Cells deposit, crosslink, and breakdown the surrounding ECM based on cues from their microenvironment. Changes to the matrix stiffness are hypothesized to trigger pathways for increased ECM deposition by activated fibroblasts, characterized by the expression of α-smooth muscle actin (α-SMA), and assumed to be essential in the development of fibrosis [25, 31]. Transcriptional activation by YAP/TAZ, coupled with higher substrate stiffness, triggers it’s localization to the nucleus which is accompanied with increased matrix production [31, 32]. The deposited matrix may subsequently undergo crosslinking and also be associated with altered MMP regulation. MMP’s are the primary enzymes linked to matrix remodeling that are associated with cancer cell migration and metastasis [33]. Increased upregulation of MMP activity in pancreatic cells in response to substrate stiffness is mediated through higher cell contractility [34]. Our results suggest that the feedback between stiffening and degradation of the ECM by HMF3S cells may hence promote early fibrosis in tissues.

### 3.4. Substrate stiffness regulates TGF-β activation

Because stiffness regulates the expression of ECM genes and MMP-2, both direct targets of TGF-β signaling, we explored for possible modulation of TGF-β expression in cells by the substrate properties of PDMS. The mRNA expression of TGF-β, obtained using qRT-PCR (Fig. 6a), shows marginal reduction at the transcript level in cells grown on PDMS substrates as compared to control dishes. These differences were however not statistically significant. We next used the CCL64-PAI assay to quantify TGF-β protein in the media. In this method, we use the promoter of PAI (plasminogen activator inhibitor) gene, a transcriptional target of TGF-β, fused with the luciferase reporter, to assay TGF-β in engineered CCL64 cells, containing stable reporters for PAI promoter constructs, that are highly sensitive to TGF-β signaling.

**Fig. 6:**
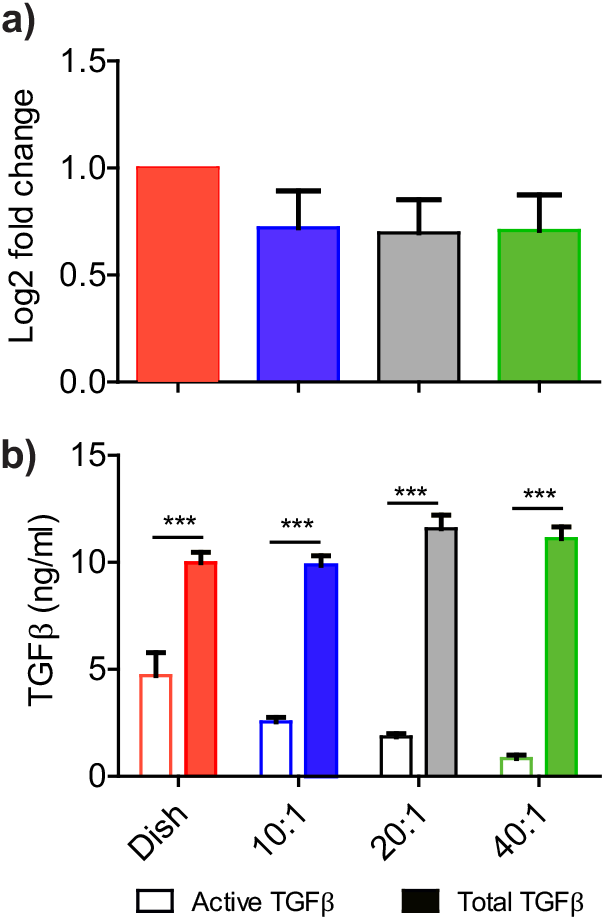
Substrate stiffness modulates TGF-β activation. (**a)** RNA collected from cells seeded on petridish, 10:1, 20:1, and 40:1 PDMS substrates were analyzed for TGF-β1 mRNA expression. (**b)** TGF-β concentration from condition media quantified using CCL64PAI assay. All experiments were repeated in triplicates and used in the reported analysis. Data are represented as mean ± SEM; *** p<0.001.

We measured the active TGF-β content in the conditioned media collected from cells cultured on different stiffness substrates. We generated the TGF-β response standard curve using recombinant TGF-β protein (Fig. S3), and next divided the serum free condition media, collected from conditioned media from cells on the different substrates, into two parts. The first was heated at 85°C for 10 minutes to activate TGF-β (active), and the second was kept on ice (native). CCL64-PAI cells were treated with the two conditioned media separately. These comparisons show that there were no differences in the heat-activated condition media from HMF3S cells on the different PDMS substrates. In contrast, there was gradual decrease in the activity of native condition media with decreasing substrate stiffness (Fig. 6b). Results hence demonstrate that the active TGF-β content in the conditioned media correlates with differential substrate stiffness.

Substrates of higher stiffness are essential for the activation of fibroblast through TGF-β which results in higher cell contractility and matrix remodeling [25]. Activation of fibroblasts into myofibroblasts alters the ECM gene expression which is regulated by TGF-β [34]. TGF-β is secreted in tissues as a latent complex consisting of latent TGF-β binding protein (LTBP), latency associated peptide (LAP), and mature TGF-β. Activation of the mature TGF-β dimer, through proteolytic cleavage of LAP and/or interaction of LTBP with integrins, is essential for receptor binding. Fibroblasts hence regulate the activation of TGF-β based on the underlying substrate stiffness. Because results also show regulation of integrin expression by the underlying substrate stiffness (Fig. 3a), we suggest that this could be a possible mechanism of TGF-β activation through changes in the substrate stiffness.

## 4. Conclusions

Form and function in cells is inextricably linked to the substrate characteristics, including stiffness. Cells use inputs from mechanical cues in the microenvironment and receive feedback *via* integrins to influence the cytoskeletal architecture and nuclear function. The specific mechanisms of how cells sense their mechanical milieu and translate these into biochemical signals are presently not well understood. Numerous challenges exist when working with natural biomaterials to control different aspects of the physical properties of the substrate; these are partially offset using well-characterized biomaterials with tuneable material properties, such as PDMS. The biocompatibility of PDMS renders it useful for application in biological studies including single cell and immunoassays, stem cell research, in microarrays and bioreactors, and to quantify cellular sensitivity to substrate topology [35-39]. The tunability of PDMS substrates makes it an attractive material in mechanobiological studies to investigate cellular responses that may be sensitive to the underlying matrix stiffness.

Our results demonstrate that substrate stiffness modulates cell morphologies and proliferations. Fibroblasts had higher spread areas on control petridishes as compared to PDMS substrates of varied stiffness. Cell nuclei were rounded on compliant substrates and correlated with increased tubulin expression levels. Cell proliferations were higher on stiffer substrates and showed cell cycle arrest on lower stiffness substrates. There was a significant decrease in the expression and signaling of integrin α5 on soft PDMS substrates as compared to stiffer ones. The expression levels of collagen I, collagen III, and MMP-2 decreased on compliant PDMS substrates as compared to stiffer ones. There were no significant changes at transcript level in MMP-9 on all substrates stiffness in this study. We used gelatin zymography to test for the differences in MMP-2 and MMP-9 activity levels from cells on varied substrates. Our data show little MMP-9 activity on all substrates, and a higher conversion of pro-MMP-2 to active-MMP-2 on lower stiffness substrates. These results show the critical feedback between substrate stiffness and ECM regulation by fibroblasts which is highly relevant in tissue fibrosis. Finally, we demonstrate a change in the activated state of the fibroblast on higher stiffness substrates that are linked to altered TGF-β activation. This study provides important insights on the relationships between substrate stiffness and cell morphologies on differential gene expressions associated with the ECM. We hope these findings are useful in better understanding of cell-matrix interactions that are critical in developing molecules to mitigate and facilitate tissue repair and fibrosis.

## 5. Author Contributions

Brijesh Kumar Verma: Conceptualization, Methodology, Investigation, Inputs on writing. Aritra Chatterjee: Methodology, Investigation, Formal analysis, Inputs on writing. Paturu Kondaiah: Conceptualization, Methodology, Reviewing, Supervision. Namrata Gundiah: Conceptualization, Methodology, Supervision, Manuscript writing, Data curation, Reviewing, Funding.

## 6. Acknowledgements

We thank the imaging facility in Biological Sciences, IISc for the confocal imaging reported in this study. PK lab is supported by IISc-DBT partnership program and DST-FIST infrastructure to MRDG. PK is a recipient of senior scientist fellowship from Indian National Science Academy. NG is grateful to the Department of Biotechnology through the Bioengineering and Biodesign Initiative for project support.

## Supplementary Figures

**Fig. S1:**
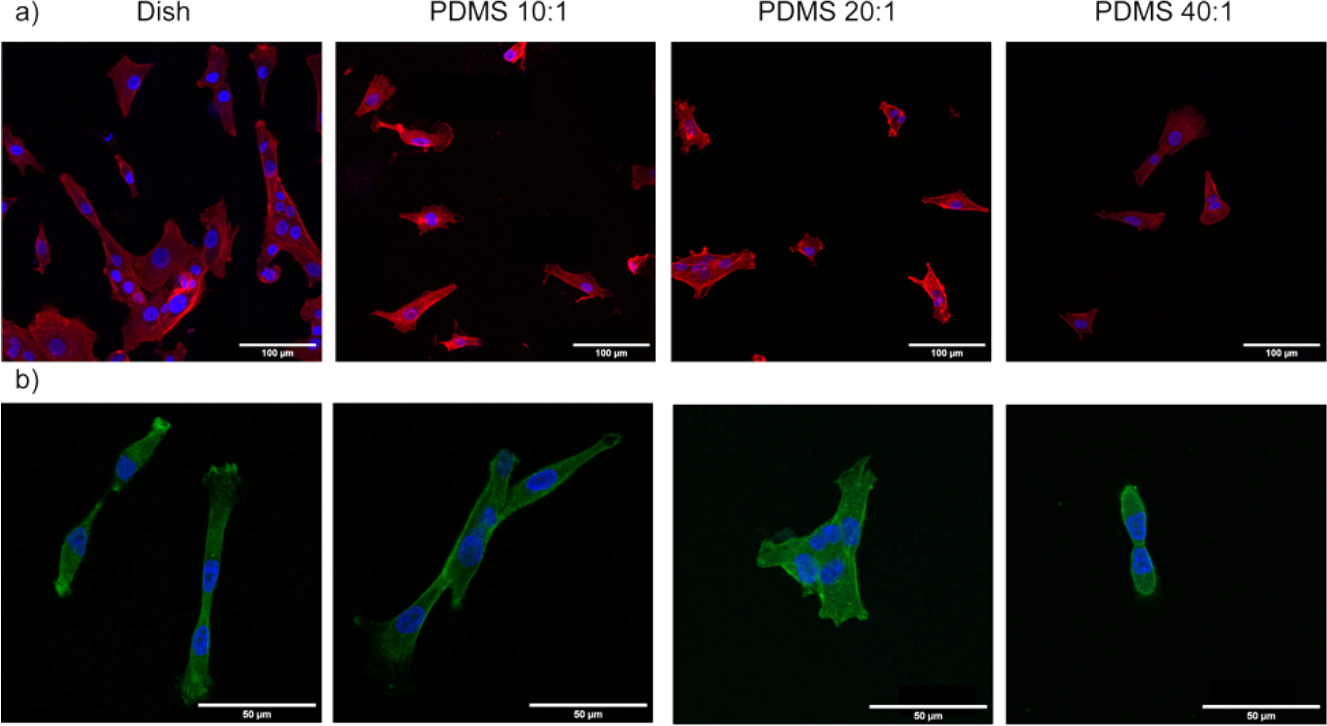
Confocal images of cells seeded on petridish, 10:1, 20:1 and 40:1 PDMS substrates show differences in the cell morphologies. **a)** The upper panel is stained for actin (red) using phalloidin red, and DAPI which stains the nuclei blue. Scale bar: 100 μm. **b)** The lower panel shows tubulin (green) and DAPI stained cells (blue). All experiments were performed in biological triplicates. Scale bar is 50 μm.

**Fig. S2:**
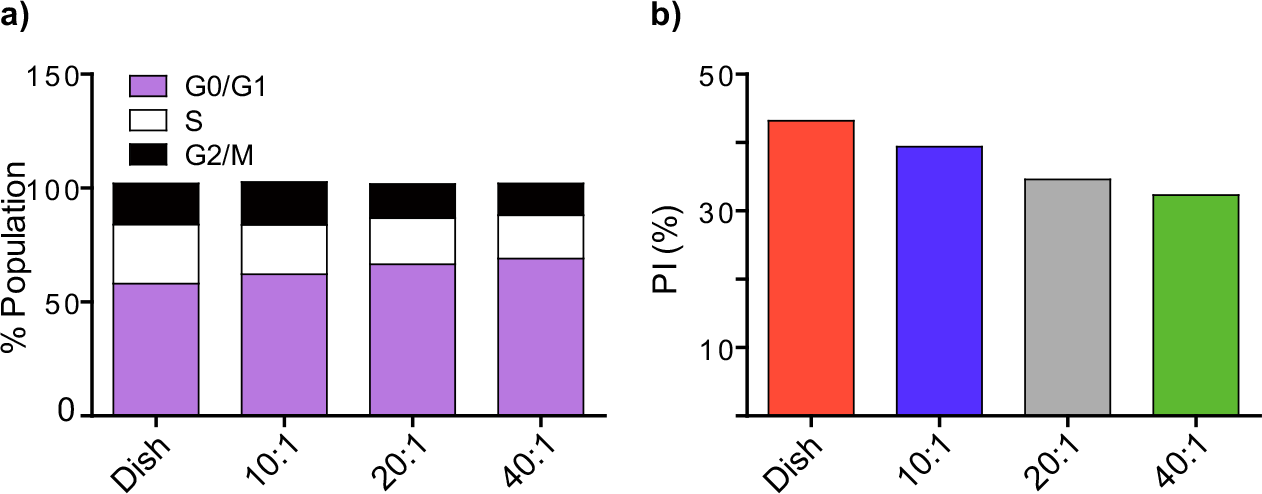
**(a)** Histogram representing the percentage of cells in different cell cycle phases shown for substrates of different stiffnesses. **(b)** The proliferative index of cells seeded on four different substrates is shown.

**Fig. S3:**
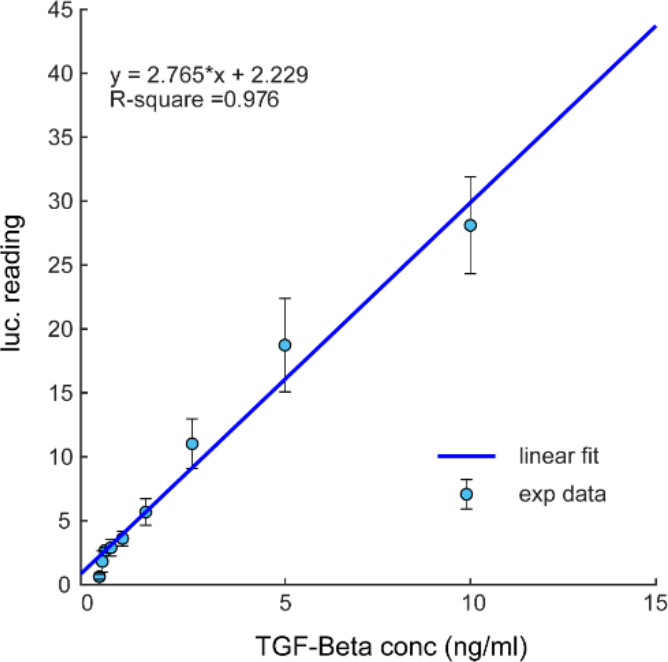
Standard curve for recombinant TGF-β protein response using CCL64 PAI assay is shown.

**Table S1:**
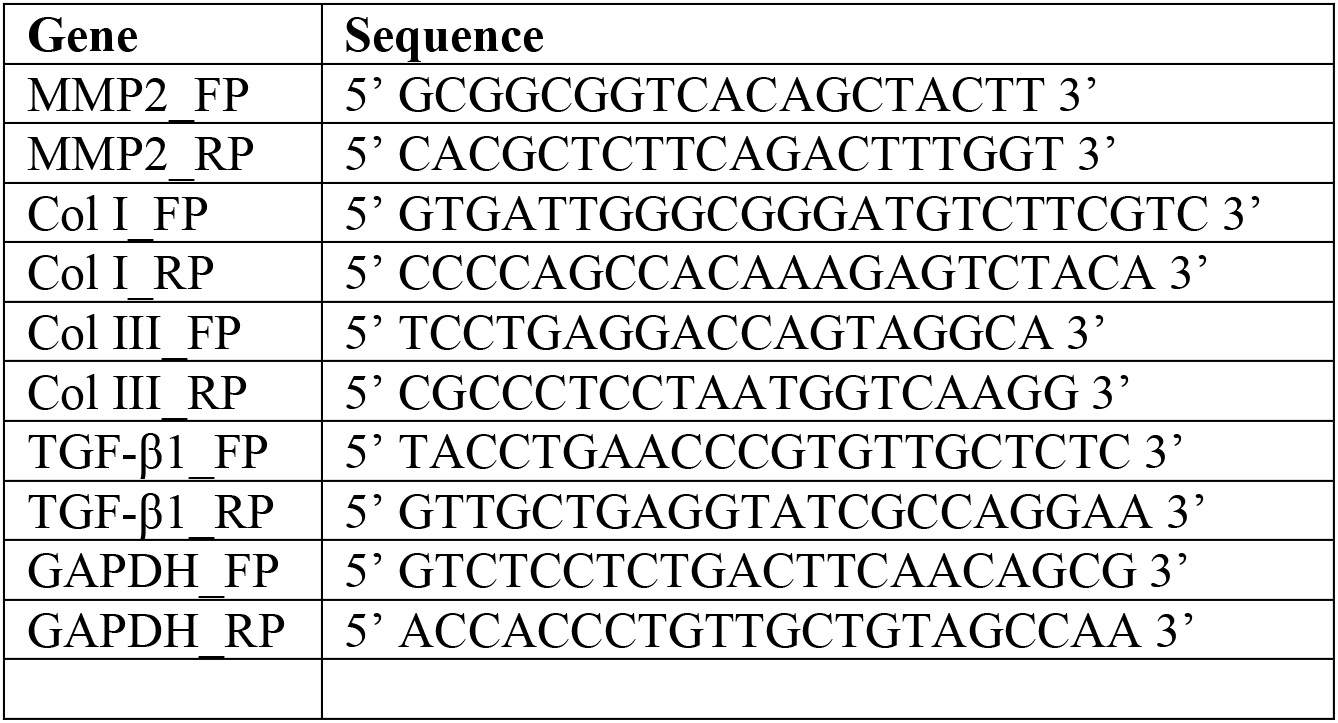
List of primers used in the study are indicated.

